# The RASSF6 tumor suppressor protein regulates apoptosis and the cell cycle progression *via* Retinoblastoma protein

**DOI:** 10.1101/334276

**Authors:** Shakhawoat Hossain, Hiroaki Iwasa, Aradhan Sarkar, Junichi Maruyama, Kyoko Arimoto-Matsuzaki, Yutaka Hata

## Abstract

RASSF6 is a member of the tumor suppressor Ras-association domain family (RASSF) proteins. *RASSF6* is frequently suppressed in human cancers and its low expression is associated with poor prognosis. RASSF6 regulates cell cycle arrest and apoptosis and plays a tumor suppressor role. Mechanistically, RASSF6 blocks MDM2-mediated p53 degradation and enhances p53 expression. However, RASSF6 also induces cell cycle arrest and apoptosis in the p53-negative background, which implies that the tumor suppressor function of RASSF6 does not depend solely on p53. In this study, we have revealed that RASSF6 mediates cell cycle arrest and apoptosis *via* pRb. RASSF6 enhances the interaction between pRb and protein phosphatase. RASSF6 also enhances *P16INK4A and P14ARF* expression through suppressing BMI1. In this way, RASSF6 increases unphosphorylated pRb and augments the interaction between pRb and E2F1. Moreover, RASSF6 induces TP73-target genes *via* pRb and E2F1 in the p53-negative background. Finally, we confirmed that RASSF6 depletion induces polypoid cells in p53-negative HCT116 cells. In conclusion, RASSF6 behaves as a tumor suppressor in cancers with the loss-of-function of p53, and pRb is implicated in this function of RASSF6.

## INTRODCUTION

Ras association domain family (RASSF) 6 is a member of the RASSF proteins (1-3). *RASSF6* is epigenetically silenced in acute lymphocytic leukemia, chronic lymphocytic leukemia, neuroblastoma, metastatic melanoma, and gastric cardia adenocarcinoma (4-8). RASSF6 suppression is more frequently observed in gastric cancer, pancreatic ductal adenocarcinoma, and gastric cardia adenocarcinoma at the advanced stage (8-10). These findings support the tumor suppressive role of RASSF6.

Exogenously expressed RASSF6 induces apoptosis in caspase-dependent and caspase-independent manners in various cells (11). Conversely RASSF6 depletion blocks tumor necrosis factor α-induced apoptosis in HeLa cells, okadaic acid-induced apoptosis in rat hepatocytes, and sorbitol-induced apoptosis in human renal proximal tubular epithelial cells (11-13). RASSF6 also causes G1/S arrest and is implicated in ultraviolet (UV)-induced cell cycle arrest (14).

The Hippo pathway is a tumor-suppressive signaling pathway that comprises mammalian Ste20-like (MST) 1/2 kinases and large tumor suppressor (LATS) 1/2 kinases (15-17). RASSF6 interacts with MST1/2 kinases and inhibits the kinase activity (12). Reciprocally MST1/2 suppress RASSF6-induced apoptosis. When cells are exposed to okadaic acid, which activates the Hippo pathway, RASSF6 and MST1/2 are dissociated. Consequently, RASSF6 induces apoptosis. In this manner RASSF6 co-operates with the Hippo pathway to function as a tumor suppressor.

RASSF6 binds to MDM2 and blocks p53 degradation by MDM2 (14). UV enhances p53 expression and triggers the transcription of p53 target genes that are implicated in apoptosis and cell cycle regulation. RASSF6 depletion attenuates UV-triggered increase of p53 expression and blocks the induction of p53 target genes. MDM2-p53 is instrumental for the tumor suppressive role of RASSF6. Nevertheless, RASSF6 induces apoptosis even in p53-compromized HeLa cells and p53-negative HCT116 (HCT116 p53-/-) cells, suggesting that RASSF6 controls apoptosis *via* a certain molecule other than p53. Modulator of apoptosis-1 (MOAP1), the activator of Bax, binds to RASSF6 (12, 18, 19). MOAP1 depletion attenuates RASSF6-induced apoptosis. The double knockdown of MOAP1 and p53, however, does not exhibit additional effect on RASSF6-induced apoptosis (14). It means that MOAP1 is placed in the same pathway as p53.

Retinoblastoma protein (pRb) and p53 are thought to be the two major tumor suppressors (20-23). *RB1* that encodes pRb is mutated in familial retinoblastoma. Unphosphorylated pRb forms a complex with E2F1 and inhibits E2F1-mediated transcription (24, 25). The phosphorylation of pRb by cyclin dependent kinases (CDKs) releases E2F1 from pRb and promotes cell cycle. In this study, we examined the implication of pRb in the tumor suppressive role of RASSF6. We have demonstrated that RASSF6 reduces the phosphorylation of pRb to enhance the interaction of pRb and E2F1. Furthermore, in the presence of RASSF6, E2F1 mediates transcription of pro-apoptotic *TP73* and *CASP7* in HCT116 p53-/- cells. Consistently, the suppression of *RB1* and *E2F1* decreases RASSF6-mediated apoptosis.

## RESULTS

### Depletion of *RB1* overrides RASSF6-indued cell cycle arrest

We tested the effect of RASSF6 on the cell cycle in the p53-negative background. Exogenously expressed RASSF6 blocked EdU incorporation in HCT116-p53-/- cells (**Fig. 1A, siCont., arrowheads)**. However, when *RB1* was knocked down (**Fig. 1C**), EdU was incorporated in RASSF6-expressing cells (**Fig. 1A, siRB1#1 and siRB1#2, arrows**). In the quantification, almost 80% cells incorporated EdU in control cells, whether *RB1* was knocked down or not (**Fig. 1B, black bars**). RASSF6 reduced the incorporation of EdU to 10% (**Fig. 1B, siCont., a gray bar**) but *RB1* silencing recovered it to about 40% **(Fig. 1B, siRB1#1 and siRB1#2, gray bars**).

**Figure 1.**
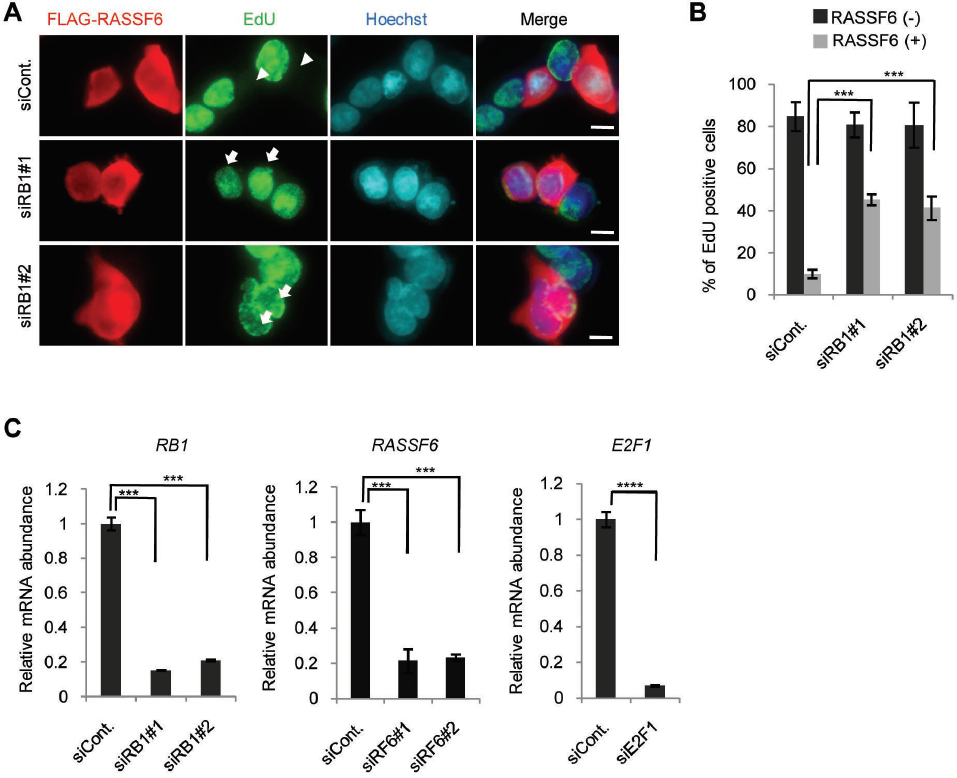
RASSF6 suppresses EdU incorporation *via* pRb. (A) HCT116 p53-/- cells were transfected with control siRNA or *RB1* siRNAs. 48 h later, the cells were replated at 5×104 cells/well in a 12-well plate and transfected with pCIneoFHF-RASSF6 (FLAG-RASSF6). 6 h after transfection, the cells were treated with 2 mM thymidine, cultured for 18 h, and then released from the thymidine block. 2 h later, the cells were treated with 10 μM EdU for 1h and fixed and immunostained with anti-FLAG-(red) and anti-EdU-(green) antibodies. The nuclei were visualized with Hoechst 33342. In the upper panel (siCont.), the arrowheads indicate that FLAG-RASSF6-expressing cells do not incorporate EdU. In the lower panels (siRB1#1 and #2), the arrows indicate FLAG-RASSF6-expressing cells that incorporate EdU. Bar, 10 μm. (B) 150 FLAG-positive cells and negative cells were observed. The ratio of the cells incorporating EdU was calculated. Three independent samples were evaluated. Data indicate the mean with S.D. ***, p<0.001. (C) Validation of *RB1, RASSF6*, and *E2F1 silencing* in HCT116 p53-/- cells. HCT116 p53-/- cells were transfected with control, two *RB1*, two *RASSF6,* and *E2F1* siRNAs. 96 h later, mRNAs were extracted and quantitative RT-PCR was performed. ***, p<0.001: and ****, p<0.0001.

### RASSF6 blocks the phosphorylation of pRb and enhances the interaction between pRb and E2F1

The interaction between pRb and E2F1 is regulated by the phosphorylation of pRb by CDKs (20, 26, 27). Phosphorylation at threonine-821 induces the intramolecular binding of the C-terminal domain to the pocket domain and blocks the interaction between pRb and E2F1 (24, 27). Phosphorylation at serine-608 also inhibits the interaction between pRb and E2F1 (28). RASSF6 remarkably reduced the phosphorylation at serine-608 and slightly attenuated the phosphorylation at threonine-821 (**Fig. 2A**). Conversely *RASSF6* silencing augmented the phosphorylation at these sites (**Fig. 2B**). Serine-807/811 are phosphorylated by CDK4 and are discussed to be required for phosphorylation at other sites (24, 29-31). Serine-807 phosphorylation is also considered to play a role in the G0/G1 transition and the binding to Bax (32, 33). RASSF6 decreased the phosphorylation at serine-807/811, while *RASSF6* silencing augmented it (**Fig. 2A and 2B**). To further confirm the effect of RASSF6, we examined the phosphorylation states of pRb in the cytoplasmic and nuclear fractions. The cytoplasmic pRb was not detected by the antibody specific for the unphosphorylated pRb, which does not recognize the phosphorylated pRb (**Fig. 2C, the third panel**). To more clearly separate phosphorylated and unphosphorylated pRb, we used Phos-tag gels, in which phosphorylated proteins migrate slowly. The analysis by Phos-tag gels revealed that the major part of the cytoplasmic pRb was phosphorylated, while most of the nuclear pRb was unphosphorylated but phosphorylated pRb was also detected in the nucleus (**Fig. 2C, an arrow, the third lane**). However, the co-expression of RASSF6 reduced the nuclear phosphorylated pRb (**Fig. 2C, an arrowhead, the sixth lane**). All these findings indicate that RASSF6 reduces the phosphorylation of pRb. In the co-immunoprecipitation experiment using HEK293FT cells, RASSF6 augmented the interaction between E2F1 and pRb, which is consistent with the decreased phosphorylation at serine-608 and threonine-821 (**Fig. 2D, an arrow**). Furthermore, RASSF6 suppressed E2F1-promoter reporter (**Fig. 2E, siCont.**), but *RB1* silencing abrogated this inhibition, supporting that RASSF6 inhibits E2F1 *via* pRb (**Fig. 2E, siRB1#1**). Consistently, *RASSF6* silencing enhanced E2F1-promoter reporter (**Fig, 2F**). It is reported that LATS2 is implicated in the assembly of DREAM, which represses E2F1 target gene transcription (34). Considering that RASSF6 cross-talks with the Hippo pathway, we silenced *LATS1* and *LATS2*. However, RASSF6 suppressed the E2F1 promoter reporter even in the *LATS1*/*2*-negatvie background (**Fig. 2G**). Moreover, the depletion of LIN52, which is a component of DREAM complex (29), did not affect RASSF6-mediated suppression of E2F1 promoter reporter (**Fig. 2H**). These findings suggest that RASSF6 functions independently of DREAM complex.

**Figure 2.**
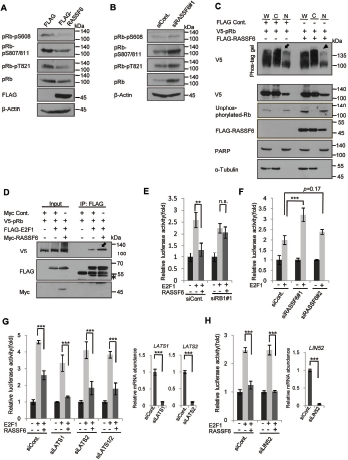
RASSF6 suppresses pRb phosphorylation and enhances the interaction between pRb and E2F1. (A) The cell lysates of parent HCT116 p53-/- cells transfected with control pCIneoFHF (FLAG) or pCIneoFHF-RASSF6 (FLAG-RASSF6) were immunoblotted with the indicated antibodies to detect endogenous pRb and to evaluate the phosphorylation.(B) HCT116 p53-/- cells were transfected with control siRNA (siCont.) or *RASSF6* siRNA (siRASSF6#1). 72 h later, the cell lysates were immunoblotted with the indicated antibodies. (C) V5-pRb (pLX304-pRb-V5) was expressed alone or with FLAG-RASSF6 in HCT116 p53-/- cells. 24 h later, the cells were harvested and the subcellular fractionation was performed. The comparable amounts of the whole cell lysates (W), the cytoplasmic fraction (C) and the nuclear fraction (N) were analyzed by SDS-PAGE for the immunoblotting with anti-FLAG, anti-V5, anti-unphosphorylated-Rb, anti-FLAG, anti-Poly (ADP-ribose) polymerase (PARP), and anti-α-tubulin antibodies, or on the Phos-tag gel for the immunoblotting with anti-V5 antibody to evaluate phosphorylated and unphosphorylated pRb. PARP and α-tubulin were used as nuclear and cytoplasmic markers. The arrow indicates the phosphorylated pRb in the nuclear fraction. As indicated by the arrowhead, RASSF6 reduced the phosphorylated pRb in the nuclear fraction. (D) HEK293FT cells were transfected with various combinations of pLX304-pRb-V5, pClneoFH-E2F1, and pClneoMyc-RASSF6. Immunoprecipitation was performed with anti-DYKDDDDK (1E6) antibody beads. The inputs and the immunoprecipitates were immunoblotted with the indicated antibodies. In the presence of RASSF6, the amount of the co-immunoprecipitated pRb was increased (the arrow). The asterisk indicates the immunoglobulin heavy chain. (E) HEK293FT cells were transfected with control siRNA or *RB1* siRNA#1. 48 h later, the cells were transfected with E2F1-Luc (−242) reporter and pCMV alkaline phosphatase. E2F1 (pCIneoFH-E2F1) and/or RASSF6 (pCIneoMyc-RASSF6) were co-expressed. 24 h later, luciferase assay was conducted by use of Picagene as a substrate. RASSF6 suppressed E2F1-mediated enhancement of luciferase activity (the second and third bars, siCont), but *RB1* knockdown abolished the effect of RASSF6. **, p<0.01; n.s., not significant. (F) HEK293FT cells were transfected with control siRNA or *RASSF6* siRNAs (#1 or #2), The reporter assay was performed as described for Figure 2E. ***, p<0.001. (G) HEK293FT cells were transfected with *LATS1* and *LATS2* siRNAs. The reporter assay was performed as described for Figure 2E. The validation of *LATS1/2* silencing was demonstrated on the right. ***, p<0.001.(H) HEK293FT cells were transfected with *LIN52* siRNA. The reporter assay was performed as described for Figure 2E. The validation of *LIN52* silencing was demonstrated on the right. ***, p<0.001.

### RASSF6 promotes the interaction between pRb and protein phosphatases

pRb phosphorylation is regulated by protein phosphatases (PP1A and PP2A) (35). In *Drosophila melanogaster*, dRASSF, a fly homologue of RASSF, interacts with fly homologues of components of the striatin-interacting protein phosphatases and kinases (STRIPAK) complexes (36). STRIPAK is a complex of PP2A that interacts with Ste20-like kinases (37). In mammals, although the direct interaction between RASSF and STRIPAK complex is not reported, RASSF proteins are detected in the interactome of MST kinases and STRIPAK complex (38). We hypothesized that RASSF6 promotes as a scaffold the interaction between pRb and protein phosphatases and facilitates dephosphorylation of pRb. We immunoprecipitated RASSF6 from human colon cancer SW480 cells and detected the co-immunoprecipitated pRb with RASSF6 (**Fig. 3A, an arrow**). To further confirm the interaction, we exogenously expressed RASSF6 and pRb in HEK293FT cells and performed the co-immunoprecipitation experiment. When FLAG-RASSF6 was immunoprecipitated, V5-pRb was co-immunoprecipitated (**Fig. 3B, the left**). In the reverse experiment, FLAG-RASSF6 was co-immunoprecipitated with V5-pRb (**Fig. 3B, the right)**. In the experiment shown in Fig 2C, we noted that RASSF6, when co-expressed with pRb, was recovered not only in the cytoplasm but also in the nucleus (**Fig. 2C, the second panel**). This observation prompted us to ask whether pRb affects the subcellular localization of RASSF6. We expressed FLAG-RASSF6 with or without V5-pRb in HeLa cells and evaluated the subcellular localization in the immunofluorescence and in the subcellular fractionation (**Fig. 3C and 3D**). FLAG-RASSF6 was distributed mainly in the cytoplasm (**Fig. 3C, the upper panel, and Fig. 3D**). However, when co-expressed with V5-pRb, FLAG-RASSF6 was detected in the nucleus (**Fig. 3C, the lower panel**). The subcellular fractionation also supported that pRb increased the nuclear RASSF6 (**Fig. 3D, an arrow**). We next examined the effect of RASSF6 on the interaction between protein phosphatases and pRb. We expressed luciferase-fused PP1A and PP2A with V5-pRb in HEK293FT cells and immunoprecipitated V5-pRb. The luciferase activity in the immunoprecipitates was measured to evaluate the co-immunoprecipitation of PP1A and PP2A with pRb. RASSF6 increased luciferase-PP1A and -PP2A co-immunoprecipitated with pRb (**Fig. 3D**). These findings are consistent with the assumption that RASSF6 promotes the interaction of pRb with protein phosphatases allowing pRb to remain unphosphorylated. RASSF5 promotes dephosphorylation of pRb (39). RASSF1A blocks dephosphorylation of mammalian Ste20-like kinases (MST1 and −2) (40). As RASSF1A and RASSF5 can interact with RASSF6 (41), it is possible that RASSF6 regulates the phosphorylation state of pRb and the interaction of pRb with E2F1 through RASSF1A and RASSF5. However, neither *RASSF1* silencing nor *RASSF5* silencing had no effect on the RASSF6-mediated suppression on E2F1 promoter reporter, which means that RASSF6 regulates pRb independently of RASSF1A and RASSF5 (**Suppl. Fig. 1**). As pRb and MDM2 interact with each other (42), we examined whether and how pRb affects the interaction between RASSF6 and MDM2. However, pRb did not show any effect (**Suppl. Fig. 2)**.

**Figure 3.**
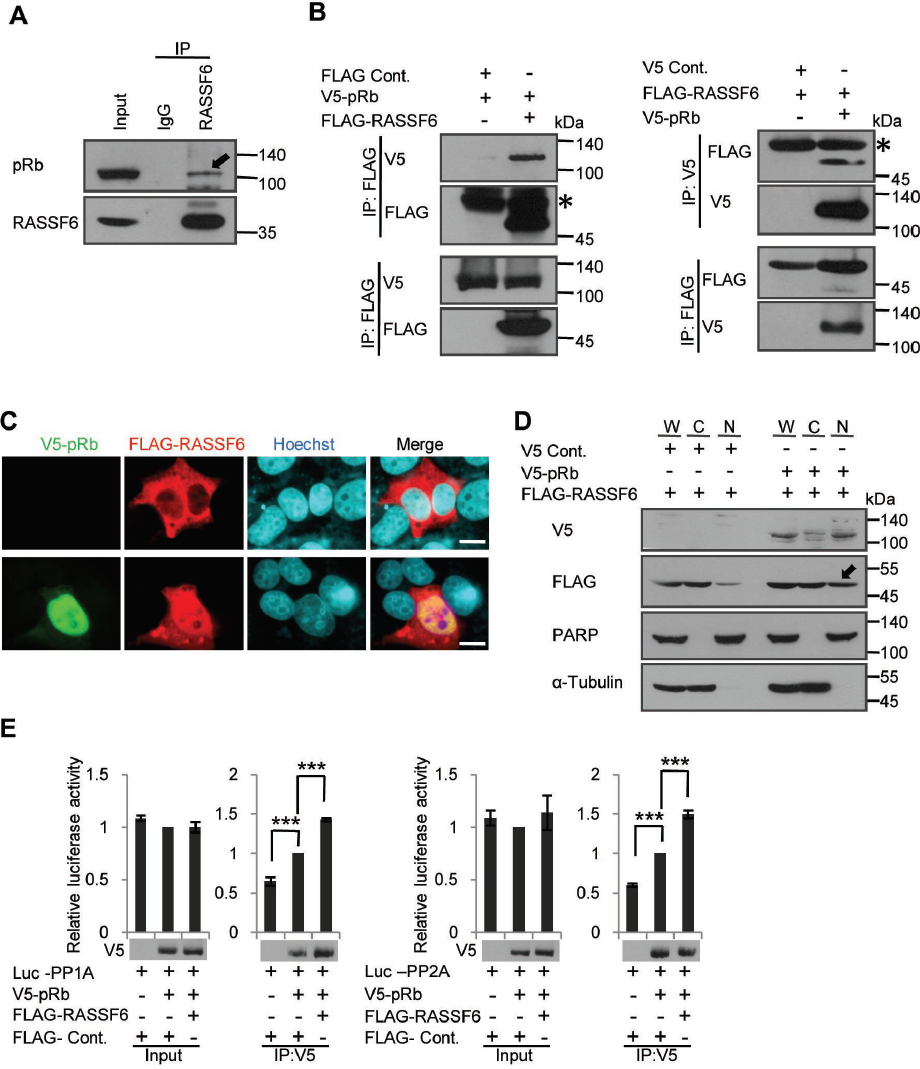
RASSF6 mediates the interaction between pRb and protein phosphatases. (A) RASSF6 was immunoprecipitated from SW480 cells. The inputs and the immunoprecipitates were immunoblotted with the indicated antibodies. The arrow indicates the co-immunoprecipitated pRb, (B) HEK293FT cells were transfected with pLX304-pRb-V5 and either control pCIneoFHF (FLAG) or pCIneoFHF-RASSF6 (FLAG-RASSF6). Immunoprecipitation was performed with anti-DYKDDDDK (1E6) antibody (the left) or anti-V5 antibody (the right) beads. The inputs and the immunoprecipitates were immunoblotted with the indicated antibodies. The asterisks indicate the immunoglobulin heavy chain. (C) and (D) FLAG-RASSF6 (pCIneoFHF-RASSF6) was expressed alone or with V5-pRb (pLX304-pRb-V5) in HeLa cells. In (C), FLAG-RASSF6 was distributed in the cytoplasm, when expressed alone, but was detected in the nucleus, when co-expressed with V5-pRb. Bar, 10 μm. In (D), the subcellular fractionation was performed. The nuclear RASSF6 was increased when co-expressed with V5-pRb (the arrow). (E) HEK293FT cells were transfected with various combinations of pClneoLuc-PP1A, pCIneoLuc-PP2A, pLX304-pRb-V5, control pCIneoFHF, and pCIneoFHF-RASSF6. The immunoprecipitation was performed with anti-V5 antibody beads. In the bottom panels, the immunoprecipitated V5-pRb was shown. Luciferase activities in the inputs and immunoprecipitates were measured by use of Picagene as a substrate. ***, p<0.001.

### RASSF6 induces *CDKN2A* independently of p53

Another explanation of the reduced phosphorylation of pRb is the inhibition of CDKs. As RASSF6 enhances p53 expression, it is reasoned that RASSF6 up-regulates *CDKN1A* (cyclin-dependent kinase inhibitor 1A) mRNA *via* p53, and inhibits the CDK2/4-mediated phosphorylation of pRb *via* CDKN1A. However, in this study, we want to know how RASSF6 functions as a tumor suppressor in the p53-negative background. We therefore examined whether RASSF6 regulates *CDKN1A* and *CDKN2A* in HCT116 p53-/- cells. The treatment with doxorubicin enhance *P16INK4A* and *P14ARF,* both of which are encoded in *CDKN2A* locus, at the mRNA level (**Fig. 4A, siCont.**). RASSF6 depletion abolished the enhancement of *P16INK4A* and *P14ARF* (**Fig. 4A, siRASSF6#1**). Doxorubicin, although to a lesser extent, enhanced *CDKN1A* in HCT116 p53-/- cells, but *RASSF6* silencing had no effect on the enhancement of *CDKN1A* (**Fig. 4A, *CDKN1A***). It implies that RASSF6 up-regulates *P16INK4A* and *P14ARF* in the p53-negative background, inhibits CDK4 *via* p16 protein, and eventually prevents the phosphorylation of pRb. Moreover, it also suggests that *CDKN1A* is regulated independently of RASSF6 in the p53-negative background.

**Figure 4.**
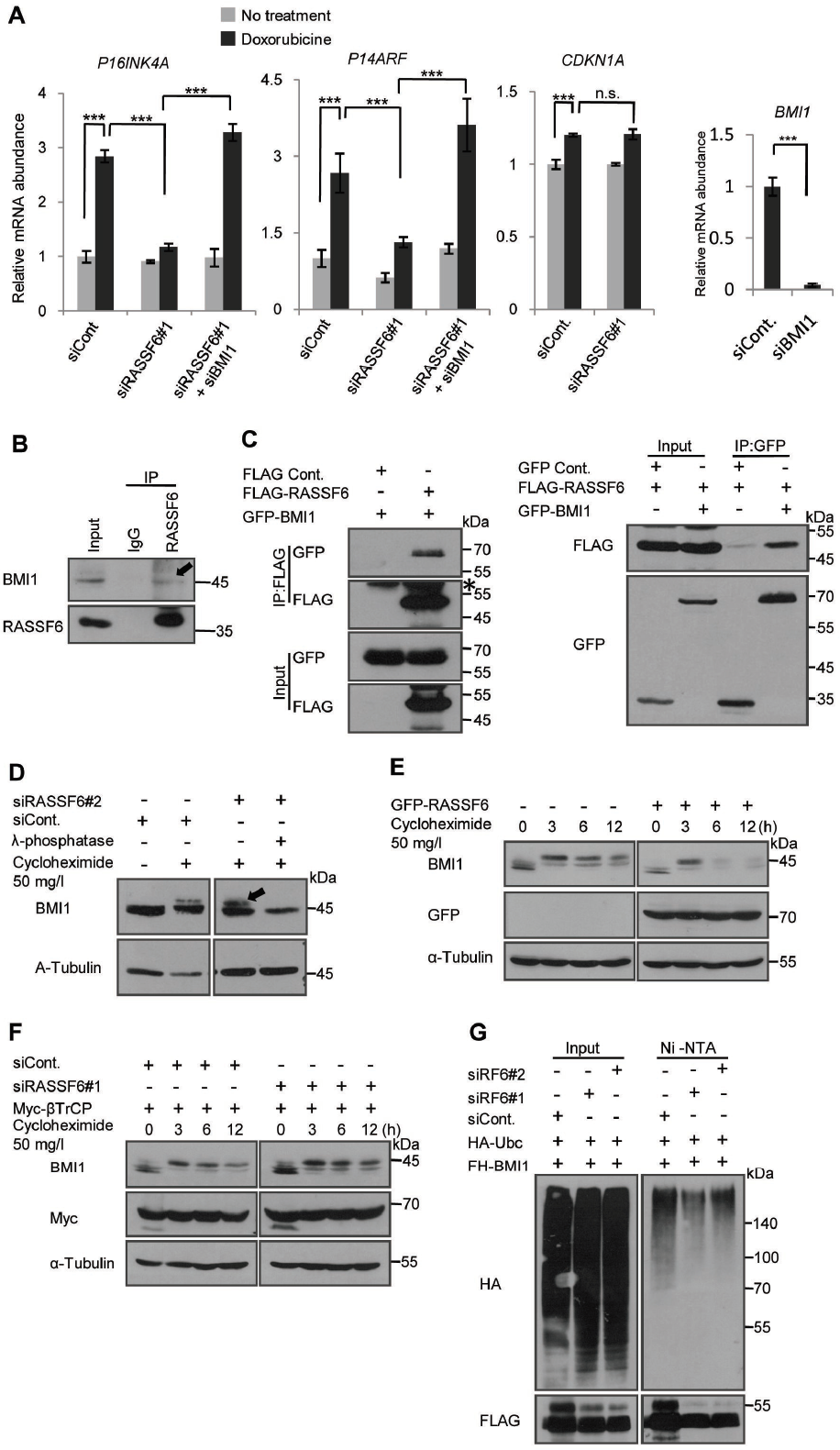
RASSF6 antagonizes BMI1 and is implicated in *P16INK4A* and *P14ARF expressions* in the p53-negative background. (A) HCT116 p53-/- cells were transfected with indicated siRNAs. 24 h later the cells were exposed to 1 mg/l doxorubicin for 1h, and then washed. The cells were further cultured and harvested to isolate mRNAs. *CDKN2A* and *P14ARF* were examined after 5 day-culture, while *CDKN1A* was evaluated after 2 day-culture with quantitative RT-PCR. *RPS18* was used as a reference. Doxorubicin enhanced *P16INK4A* and *P14ARF* expressions. Doxorubicin also enhanced *CDKN1A* expression, but to a lesser extent. *RASSF6* knockdown abolished the effect of doxorubicin. However, the additional *BMI1* silencing recovered doxorubicin-induced enhancement of *P16INK4A* and *P14ARF*. **, p<0.01; ***, p<0.001; and n.s., not significant. The validation of *BMI1* silencing was performed by qRT-PCR. (B) RASSF6 was immunoprecipitated from SW480 cells. The inputs and the immunoprecipitates were immunoblotted with the indicated antibodies. The arrow indicates the co-immunoprecipitated BMI1. (C) HEK293FT cells were transfected with control pCIneoFHF (FLAG), pCIneoFHF-RASSF6 (FLAG-RASSF6), control pCIneoGFP (GFP), and pCIneoGFP-BMI1 (GFP-BMI1) as indicated. The immunoprecipitation was performed with either anti-DYKDDDDK (1E6) antibody beads (the right) or anti-GFP antibody fixed on protein G sepharose fast flow 4 beads (the left). The asterisk indicates the immunoglobulin heavy chain. (D) HCT116 p53-/- cells were transfected with control siRNA (siCont.) or *RASSF6* siRNA (siRASSF6#2). 72 h later, the cells were treated with 50 mg/l cycloheximide for 3 h. The treatment with lambda phosphatase abolished the upper band. The samples were immunoblotted with the indicated antibodies. Endogenous BMI1 was detected. (E) HCT116 p53-/- cells were transfected with pCIneoGFP-RASSF6. 24 h later, the cells were treated with 50 mg/l cycloheximide and collected at the indicated time points. Endogenous BMI1 was immunoblotted. RASSF6 reduced phosphorylated BMI1 and facilitated BMI degradation. (F) HCT116 p53-/- cells were transfected with control siRNA (siCont.) or *RASSF6* siRNA (siRASSF6#1). 24 h later, the cells were transfected with pcDNA3-myc3-β TrCP. 48 h later, BMI1 degradation was evaluated as described for Figure 5D. *RASSF6* silencing increased phosphorylated BMI1 and delayed BMI1 degradation. (G) HEK293FT cells were transfected with control siRNA or *RASSF6* siRNAs (#1 and #2). 24 h later, the cells were transfected with pCIneFH-BMI1 and pCGN-HA-UBC. 48 h later, the cells were treated with 30 μ M MG-132 for 6 h and then the cells were lysed with the buffer containing guanidine hydrochloride. FH-BMI1 was isolated with NiNTA beads and immunoblotted with anti-HA antibody. *RASSF6* silencing reduced BMI1 ubiquitination.

### RASSF6 antagonizes BMI1

The next question is the mechanism by which RASSF6 enhances *P16INK4A* and *P14ARF*. As polycomb complex protein BMI1 is well-known to down-regulate *P16INK4A* and *P14ARF* expression, we suspected the implication of BMI1 (43). We examined whether *BMI1* silencing antagonized *RASSF6* silencing. As expected, the additional knockdown of *BMI1* recovered the enhancement of *P16INK4A* and *P14ARF* in RASSF6-depleted cells (**Fig. 4A**, **siRASSF6#1+siBMI1**). This observation supports that RASSF6 antagonizes BMI1 in the regulation of *P16INK4A* and *P14ARF*. RASSF6 expression did not affect *BMI1* mRNA expression (**data not shown**). We next tested whether and how RASSF6 affects BMI1 at the protein level. First, we tested the interaction between RASSF6 and BMI1. BMI1 was co-immunoprecipitated with RASSF6 from SW480 cells (Figure 4B, an arrow). We could also detect the interaction between exogenously expressed RASSF6 and BMI1 (**Fig. 4C, the right and the left**). We next examined the effect of RASSF6 on the stability of BMI1. We first confirmed that the cycloheximide treatment induced the mobility shift of BMI1 as previously reported (**Fig. 4D, the first and second lanes**) (44). *RASSF6* silencing increased the upward shifted band (**Fig. 4D, the third lane, an arrow**). The treatment with λ phosphatase abolished this shift (**Fig. 4D, the fourth lane**). These findings support that cycloheximide induces the phosphorylation of BMI1 and that *RASSF6* silencing increases it. We next evaluated BMI1 degradation in the presence of RASSF6 (**Fig. 4E**). GFP-RASSF6 slightly reduced BMI1 protein expression (**Fig. 4E, 0 h)**. More remarkably, RASSF6 decreased the upward-shifted BMI1 (**Fig. 4E, 3 h to 12 h**). βTrCP regulates BMI1 ubiquitination and degradation (44). We next examined the effect of *RASSF6* silencing on BMI1 stability in the presence of βTrCP. *RASSF6* silencing increased the up-ward shifted BMI1 and delayed its degradation (**Fig. 4F, the right**). Moreover, *RASSF6* silencing reduced BMI1 polyubiquitination (**Fig. 4G)**. We also examined the subcellular distribution of RASSF6 and BMI1 (**Suppl. Fig. 3**). RASSF6 did not affect the subcellular localization, but reduced BMI1 expression. BMI1 did not affect the subcellular localization of RASSF6, either. All these findings support that RASSF6 promotes the degradation of BMI1, which may contribute the enhancement of *P16INK4A* and *P14ARF*.

### pRb is involved in RASSF6-induced apoptosis

In certain circumstances, pRb is involved in apoptosis (45). For instance, in prostate cancer cells, cell detachment increases unphosphorylated pRb and induces apoptosis (46). Moreover, under DNA damage, pRb is dephosphorylated at CDK-mediated phosphorylation sites but phosphorylated by checkpoint kinases (47). Phosphorylated pRb binds to E2F1 and enhances pro-apoptotic genes (47). We first raised a question whether pRb is involved in RASSF6-induced apoptosis. We expressed GFP-RASSF6 in HCT116 p53-/- cells. GFP-RASSF6 caused nuclear condensation and induced cytochrome-C release (**Fig. 5A, siCont., GFP-RASSF6, arrows**). *RB1* silencing decreased nuclear condensation and cytochrome-C release (**Fig. 5A, siRB1#1 and siRB1#2, GFP-RASSF6, arrowheads**). In DNA content analysis by flow cytometry, GFP-RASSF6 increased sub-G1 population (**Fig. 5B, siCont., GFP and GFP-RASSF6**). *RB1* silencing, however, reduced sub-G1 population in RASSF6-expressing cells (**Fig. 5B, siCont and siRB1#1, GFP-RASSF6**). These findings support that pRb is involved in RASSF6-induced apoptosis in the p53-negative background.

**Figure 5.**
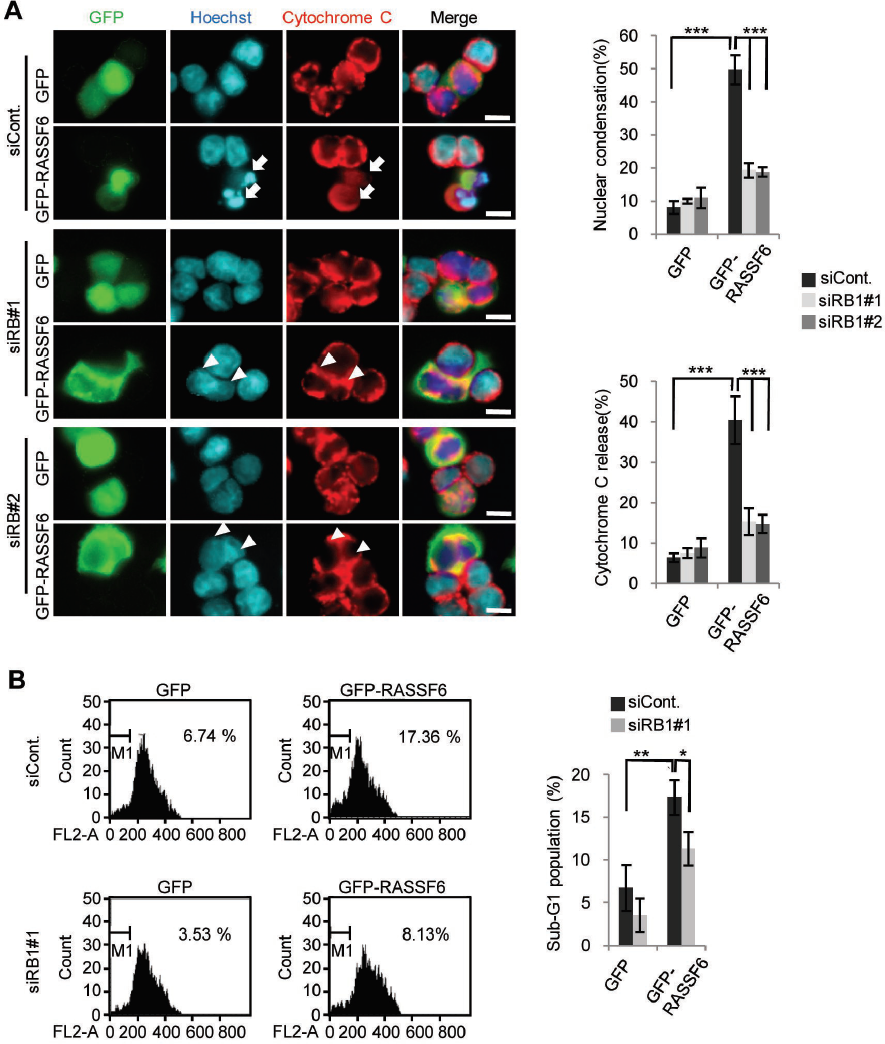
pRb is implicated in RASSF6-induced apoptosis in HCT116 p53-/- cells. HCT116 p53-/- cells were transfected with control siRNA or *RB1* siRNAs. 48 h later, the cells were replated on cover slips and were transfected with control pCIneoGFP (GFP) or pCIneoGFP-RASSF6 (GFP-RASSF6). In (A), 24 h later, the cells were immunostained with anti-cytochrome-C antibodies. The nuclei were visualized with Hoechst 33342. 50 GFP-positive cells were observed and the ratios of the cells with nuclear condensation and cytochrome-C release were calculated. GFP-RASSF6 induced nuclear condensation (arrows) but *RB1* silencing blocked it (arrowheads). Data indicate the mean with S.D. ***, p<0.001. Scale bars, 10 μ m. In (B), the cells were fixed in ice-cold 70 %(v/v) ethanol, washed with PBS, and resuspended in PBS containing 10 mg/l propidium iodide and 1 g/l RNaseA. The sub-G1 population was evaluated with FACS Calibur (BD Biosciences). The data were analyzed by use of BD CellQuest Pro Software. Data indicate the mean with S.D. *, p<0.05; **, p<0.01.

### E2F1 is involved in RASSF6-induced apoptosis

E2F1 regulates a wide variety of genes involved in not only proliferation but also apoptosis (48). For instance, E2F1 regulates apoptosis-related genes such as *APAF1* and *CASP*s (49, 50). pRb recruits histone deacetylases to repress classic E2F1 target genes such as *CCNA2*, but in cells exposed to DNA damage, pRb forms a transcriptionally active complex including E2F1 and P/CAF to promote transcription of pro-apoptotic genes such as *TP73* and *CASP7* (47, 51, 52). To determine whether E2F1 is involved in RASSF6-induced apoptosis, we examined the effect of *E2F1* silencing. *E2F1* silencing attenuated RASSF6-induced nuclear condensation and cytochrome-C release in RASSF6-expressing HCT116 p53-/- cells (**Fig. 6A**). To more directly confirm that RASSF6 plays a role in pRb/E2F1-mediated transcription of pro-apoptotic genes, we quantified mRNAs of *TP73, CASP7, and BAX*. RASSF6 increased pro-apoptotic *TP73, CASP7*, and *BAX*, but had no effect on*CCNA2*. **(Fig. 6B, siCont, gray and black bars**). However, *RB1* silencing and *E2F1* silencing abolished the effect of RASSF6 (**Fig. 6B, siRB1#1 and siE2F1**). These findings support that E2F1 is involved in RASSF6-induced apoptosis. Although YAP1, a target of the Hippo pathway, co-operates with p73, *YAP1* silencing had no effect on RASSF6-mediated apoptosis in HCT116 p53-/- cells (**Suppl. Fig. 4)**. *BMI1* silencing did not augment RASSF6-indced apoptosis, either (**Suppl. Fig. 5)**.

**Figure 6.**
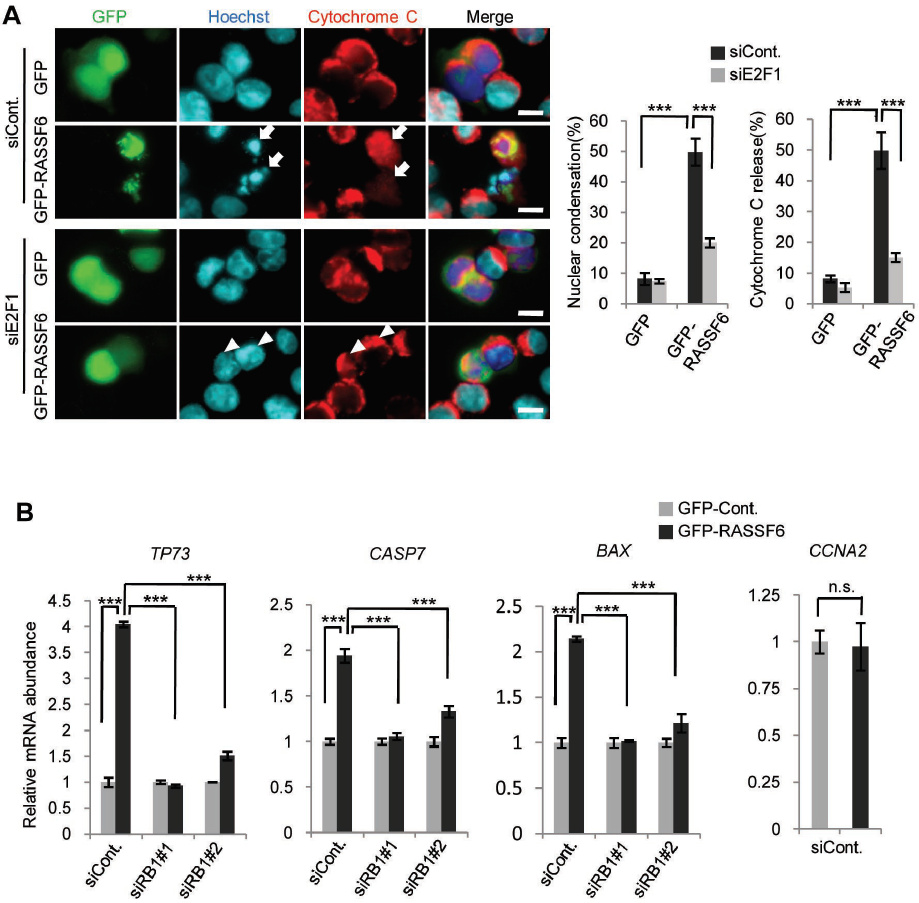
E2F1 is implicated in RASSF6-induced apoptosis in HCT116 p53-/- cells. (A) HCT116 p53-/- cells were transfected with control siRNA or *E2F1* siRNA. 48 h later, the cells were replated on cover slips and transfected with control pCIneoGFP (GFP) or pCIneoGFP-RASSF6 (GFP-RASSF6). Apoptosis was evaluated as described for Figure 5A. Data indicate the mean with S.D. ***, p<0.001. Scale bars, 10 μ m. (B) HCT116 p53-/- cells were transfected with control siRNA, *RB1* siRNA#1, or *E2F1* siRNA. 48 h later, the cells were transfected with control pCIneoGFP or pCIneoGFP-RASSF6. 24 h later, the cells were harvested and mRNAs were collected. qRT-PCR was performed by use of glyceraldehyde-3-phosphate dehydrogenase as a reference. The value for the cells expressing control GFP was set at 1.0. Data indicate the mean with S.D. for the triplicate samples. *TP73, CASP7*, and *BAX* genes were up-regulated by RASSF6. *RB1* or *E2F1* knockdown abolished the effect of RASSF6. *CCNA2* gene was not enhanced by RASSF6. ***, p<0.001; and n.s., not significant.

### RASSF6 depletion impairs DNA repair and causes genomic instability

In the previous study, we demonstrated that RASSF6 depletion impairs DNA repair in HCT116 cells and results in the generation of polyploid cells (14). To evaluate the significance of RASSF6 as a tumor suppressor in the p53-negative background, we examined the effect of RASSF6 depletion on DNA repair in HCT116 p53-/- cells. After 3 h-treatment with VP-16, γH2A.X signals appeared (**Fig. 7A, HCT116 p53+/+, 0 h**) but disappeared within 3 h after the removal of VP-16 in HCT116 p53+/+ cells (**Fig. 7A, HCT116 p53+/+, 3 h**). However, in HCT116 p53-/- cells, the signals remained up to 9 h (**Fig. 7A, HCT116 p53-/-, siCont, 9 h**). This means that DNA repair is delayed in HCT116 p53-/- cells. Even so, γH2A.X signals decreased in the time-dependent manner and became barely detectable 21 h later (**Fig. 7A, HCT116 p53-/-, siCont, 21 h**). However, when RASSF6 was suppressed, γH2A.X signals were still visible at 21 h (**Fig. 7A, HCT116 p53-/-, siRASSF6#1, 21 h**). These findings support that RASSF6 is necessary for DNA repair in the p53-negative background. We cultured VP-16-treated HCT116 p53-/- cells for 96 h and analyzed DNA content by FACS. The polyploid cells were increased by the treatment with VP-16 (**Fig. 7B, siCont, 1.64 % vs 13.03 %**). RASSF6 depletion further increased the polyploid cells (**Fig. 7B, VP-16, siRASSF6#1 and siRASSF6#2**). All these findings support the possibility that RASSF6 plays a tumor suppressor role in cancer cells with dysregulated p53

**Figure 7.**
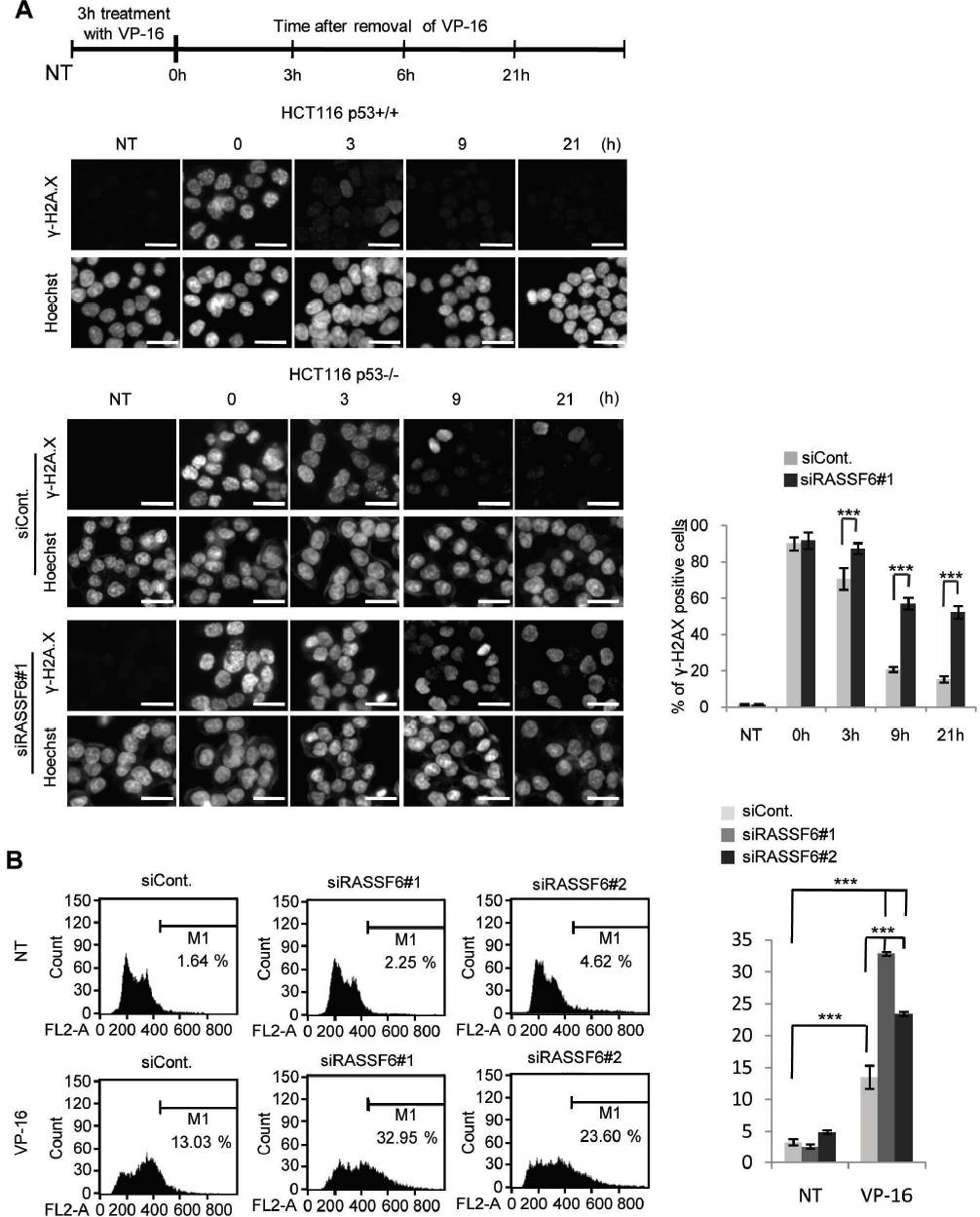
RASSF6 depletion delays DNA repair and increases polyploid cells in the p53-negative background. HCT116 p53+/+ cells were exposed to 50 μ M VP-16 for 3 h and then VP-16 was removed. The cells were harvested at the indicated time points and γH2A.X was immunostained. The scheme of the protocol was demonstrated on the top in (A). HCT116 p53-/- cells were transfected with control siRNA or *RASSF6* siRNA (#1 in (A); #1 and #2 in (B)). 48 h later, the cells were exposed to 50 μ M VP-16 for 3 h and then VP-16 was removed. In (A), thereafter, γ H2A.X was immunostained at the indicated time points (0 h means “immediately after 3 h-treatment with VP-16”) and γH2A.X was immunostained. 500 cells were observed for each sample and γH2A.X-positive cells were counted. The bargraphs are the summary of three independent experiments. Data indicate the mean with S.D. ***, p<0.001. Scale bars, 20 μ m. In (B), the cells were cultured for 96 h after removal of VP-16, and DNA contents were evaluated by use of FACS. VP-16 treatment induced polyploid cells, and RASSF6 depletion further increased them.

## DISCUSSION

RASSF6 is one of the classical RASSF proteins, the proteins that have the Ras-association domains in the C-terminal region, and co-operates with the well-conserved tumor suppressor Hippo pathway (41). The low expression of RASSF6 is observed in human cancers and correlates with the shortened disease-free survival, which corroborates that RASSF6 plays a tumor suppressive role in human. Forced expression of RASSF6 induces apoptosis and cell cycle arrest in various cells (11, 12). p53 depletion attenuates RASSF6-induced apoptosis and overcomes RASSF6-induced cell cycle arrest, which supports that p53 functions down-stream of RASSF6 (14). In fact, RASSF6 interacts with MDM2 and blocks MDM2-mediated degradation of p53. RASSF6 depletion suppresses ultraviolet exposure-induced p53 target gene expression. KRAS strengthens the interaction between RASSF6 and MDM2, and causes apoptosis depending on p53 (19). In this respect, MDM2-p53 is important for the tumor suppressor role of RASSF6. However, p53 depletion does not completely abolish RASSF6-mediated apoptosis (14). Moreover, RASSF6 induces apoptosis in p53-negative HCT116 cells. These findings imply that RASSF6 causes apoptosis independently of p53. This observation is important, because it means that RASSF6 works as a tumor suppressor in cancers with p53 mutations. In this line, we have studied the molecular mechanism underlying the p53-independent tumor suppressor role of RASSF6.

*RB1* is the first identified tumor suppressor gene and its defects cause many human cancers (23). A recent report has revealed that Nore1A (a splicing variant of RASSF5) forms a complex composed of protein phosphatase 1A and pRb to promote pRb dephosphorylation and to mediate Ras-induced senescence (39). Based on these data, we examined the relations of RASSF6 and pRb. To accentuate the role of pRb, we used HCT116 p53-/- cells in this study. First, we confirmed that *RB1* silencing attenuates RASSF6-induced cell cycle arrest (**Fig. 1**).

Unphosphorylated pRb binds and inhibits the E2F family transcription factors (20, 26). When pRb is phosphorylated by CDKs, it fails to bind the E2F proteins, so that cells proceed from G1 to enter S. During M to G1, pRb is dephosphorylated by protein phosphatases and returns to the unphosphorylated form (35). As RASSF6 blocks the DNA synthesis depending on pRb (**Fig. 1**) and Nore1A promotes pRb dephosphorylation (39), we suspected that RASSF6 as well suppresses the phosphorylation of pRb. As expected, RASSF6 reduces the phosphorylation at serine-608 and at threonine-821, both of which destabilize the interaction between pRb and E2F (**Fig. 2**). Accordingly, RASSF6 strengthens the binding of pRb to E2F1 and suppresses E2F1 promoter reporter depending on pRb (**Fig. 2D and 2E**). Conversely, *RASSF6* silencing enhanced E2F1 promoter reporter (**Fig. 2F**). A kinase shRNA screening revealed that LATS2 promotes the assembly of DREAM repressor complex and suppresses E2Ftarget genes (34). As RASSF6 cross-talks with the Hippo pathway, it is reasonable to question whether LATS2 is implicated in RASSF6-mediated suppression of E2F1 reporter activity. However, *LATS1*/*2* silencing had no effect (**Fig. 2G**). Moreover, *LIN52* silencing did not affect RASSF6-mediated repression, either (**Fig. 2H**). These findings suggest that RASSF6 repress E2F1-mediated transcription independently of DREAM repressor complex. Likewise, *YAP1* silencing did not show any effect on RASSF6-mediated apoptosis (**Suppl. Fig. 4**). This result is comprehensible, because RASSF6 plays a tumor suppressor role independently of the Hippo pathway (12).

As reported for Nore1A, RASSF6 increases the interaction between pRb and protein phosphatases. RASSF6 and pRb were co-immunoprecipitated (**Fig. 3A and 3B**). pRb recruited RASSF6 into the nucleus (**Fig. 3C and 3D**). These findings support that RASSF6 and pRb interact with each other. Considering that proteomic analysis revealed RASSF proteins in the interactome of MST kinases and STRIPAK complex, we speculated that RASSF6 may link protein phosphatases to pRb. Indeed, RASSF6 promoted the binding of PP1A and PP2A to pRb (**Fig. 3E**). RASSF6 interacts with other RASSF proteins (41). RASSF5 promotes dephosphorylation of pRb *via* PP1A (39). RASSF1A conversely blocks PP1- and PP2A-mediated dephosphorylation of mammalian Ste20-like kinase 1 and −2 (40). Therefore, it is possible that RASSF6 modulates phosphorylation of pRb *via* RASSF1A and RASSF5. However, *RASSF1* silencing or *RASSF5* silencing did not affect the inhibitory effect of RASSF6 on E1F1 promoter reporter, which indicates that RASSF6 regulates pRb independently of RASSF1 or RASSF5 (**Suppl. Fig 1**).

We also found that RASSF6 increases *P16INK4A* and *P14ARF* expression in the p53-negative background. RASSF6 promotes the degradation of BMI1, which suppresses *P16INK4A* and *P14ARF*. Serine 110 phosphorylation is known to stabilize BMI1 (53). It is considered that RASSF6 promotes dephosphorylation of BMI1 and destabilizes it. We also observed that *CDKN1A* expression was enhanced by doxorubicin in HCT116 p53-/- cells, which is consistent with a previous report (54) but that *RASSF6* silencing had no effect on *CDKN1A*. In that paper, the researchers demonstrated the implication of p63. Thus, the induction of *CDKN1A* by doxorubicin in HCT116 p53-/- cells may be independent of RASSF6. In short, RASSF6 reduces the phosphorylation of pRb in two ways; through the promotion of dephosphorylation and the inhibition of phosphorylation by CDKs (**Fig 8**).

**Figure 8.**
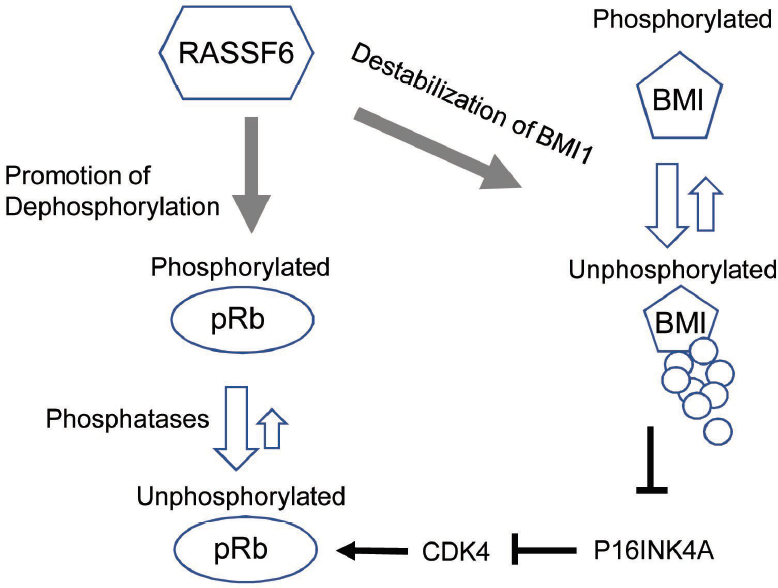
The mechanism by which RASSF6 keeps pRb unphosphorylated. RASSF6 links protein phosphatases to pRb and promotes dephosphorylation of pRb. RASSF6 also reduces the phosphorylated form of BMI1, which is resistant against βTrCP-mediated degradation, eventually destabilizes BMI1, and increases P16INK4A, which in turn inhibits CDK4.

In cells exposed to DNA damage such as doxorubicin treatment, phosphorylated pRb forms a complex with E2F1 (47). DNA damage triggers the binding of histone acetyltransferase to E2F1 and the acetylation of E2F1 and promotes the association of E2F1 with the promoters of proapoptotic genes. Therefore, we examined whether pRb and E2F1 are implicated in RASSF6-induced apoptosis. The silencing of *RB1* or *E2F1* suppresses RASSF6-induced apoptosis (**Fig. 5 and Fig. 6**). RASSF6 expression enhances proapoptotic genes including *TP73, CASP7*, and *BAX*, while the silencing of *RB1* or *E2F1* blocks the effect of RASSF6 (**Fig. 6)**. We previously observed that *TP73* silencing did not attenuate RASSF6-induced apoptosis in HCT116 p53+/+ cells (14). Therefore, the enhancement of *TP73* by RASSF6 was unexpected. However, it is discussed that p73 functionally replaces p53 in p53-deficient cells (55). As we used HCT116 p53-/- cells in this study, we speculate that the transcriptional regulatory networks may be changed by p53 depletion in these cells.

In the final set of the experiments, we showed that RASSF6 depletion impairs DNA repair in the p53-negative cells (**Fig. 7**). RASSF6 depletion also increases the generation of polyploid cells. These findings suggest that RASSF6 is a significant tumor suppressor in p53-compromized cells. Of note, *RB1* silencing does not completely abolish RASSF6-induced apoptosis in p53-negative cells (**Fig. 5**). RASSF6 induces apoptosis and suppresses proliferation in Saos-2 cells, which have *TP53* mutation and *RB1* mutation (**data not shown**). These findings mean that there should be a certain mechanism by which RASSF6 works as a tumor suppressor independently of both p53 and pRb. The dissection of such a mechanism will also be the subject of the next project.

## MATERIALS AND METHODS

### DNA constructions and virus production

pCIneoGFP, pCIneoFH, pCIneoMyc, pCIneoFHF-RASSF6, pCIneoMyc-RASSF6, pCIneoGFP-RASSF6, pCIneoLuc-PP1A, pCIneoLuc-PP2A, pLenti-EF-ires-blast, and pCIneoHAHA-MDM2 are described previously (11, 12, 14, 56). pLX304-pRb-V5 was derived from CCSB-Broad Lentiviral Expression Library (GE Healthcare Dharmacon Inc.). NheI/SalI fragment from pCIneoFHF-RASSF6 was ligated into SpeI/XhoI sites of pLenti-EF-ires-blast to generate pLenti-EF-FHF-RASSF6-ires-blast. pcDNA E2F1 is a gift of Masa-Aki Ikeda (Tokyo Medical and Dental University). PCR was performed with primers (H3086, 5’-gaattcatggccttggccggggcc-3’ and H3087, 5’-gatatcagaaatccaggggggtgag-3’) on pcDNA E2F1 and the product was digested with EcoRI/EcoRV and ligated into EcoRI/SmaI sites of pCIneoFH to generate pCIneoFH-E2F1. pCGN-HA-Ubc is a gift of Akira Kikuchi (Osaka University). Human BMI1 cDNA was obtained by PCR using the primers (H-2539, 5’-acgcgtatgcatcgaacaacgagaat-3’ and H-2540, 5’-gtcgactcaaccagaagaagttgctg-3’) and human lung and kidney cDNA libraries as the template. The PCR product was cloned into MluI/Sall sites of pCIneoGFP to generate pCIneoGFP-BMI1. pcDNA3-myc3-βTrCP was a gift of Yue Xiong (Addgene plasmid #20718) (57).

### Antibodies and reagents

The antibodies and the reagents were obtained from commercial sources: rabbit anti-GFP (598) and rat anti-HA (561) (Medical and Biological Laboratories Co. Ltd., Nagoya, Japan); mouse anti-β-actin (A1978); pepstatin A (P5318), and Hoechst 33342 (Sigma-Aldrich, St. Louis, Dallas, USA); anti-DYKDDDDK-tag (014-22383), anti-DYKDDDDK-tag beads (016-22784), anti-V5-tag-beads (016-24381), and leupeptin (334-40414) (Wako Pure Chemical Industries, Osaka, Japan); MG-132 (Nakalai tesque, Tokyo, Japan); lambda phosphatase (sc-200312) (Santa Cruz Biotechnology, Dallas, TX, USA)): rabbit polyclonal anti-V5-tag (PM003) (Medical & Biological Laboratories Co. Ltd., Nagoya, Japan); rabbit anti-RASSF6 (11921-1-AP) (Proteintech, Rosemond, Illinois, USA); protein G sepharose 4 fast flow (GE Healthcare, Little Chalfont, United Kingdom); mouse anti-cytochrome C (6H2 B4) (556432) and mouse anti-human Rb (554136), and mouse anti-underphosphorylated-Rb (554164) (BD Biosciences, San Jose, California, USA); mouse anti-Myc (9E10) (American Type Culture Collection, Manassas, Virginia, USA); rabbit polyclonal anti-phospho-Rb (Ser608) (2181), rabbit polyclonal anti-phospho-Rb (Ser807/811) (9308) (Cell Signaling Technology, Danvers, Massachusetts, USA); mouse monoclonal anti-Bmi-1 antibody (05-637) (Merck, Kenilworth, New Jersey, USA); and rabbit polyclonal anti-phospho-Rb (Thr821) (44-582G) (Thermo Fisher Scientific, Waltham, Massachusetts, USA).

### Cell cultures, transfection, and infection

HEK293FT, HCT116, HCT116 p53-/-, and HeLa cells were cultured in Dulbecco’s Modified Eagle Medium containing 10%(v/v) fetal bovine serum and 10 mM Hepes-NaOH pH7.4 under 5% CO2 at 37°C. Transfection was performed with Lipofectamine 2000 (Thermo Fisher Scientific, Waltham, USA). HEK293FT cells were transfected with pLenti-EF-FHF-RASSF6 and packaging plasmids to generate the lentivirus vector.

### Immunoprecipitation of RASSF6 from SW480 cells

SW480 cells at 50-60% in one 100-mm dish were suspended in 1 ml of the phosphate buffer saline (PBS) supplemented with 50 μM p-(amidinophenyl)methanesulfonyl fluoride (APMSF), 10 mg/l leupeptin, 3 mg/l pepstatin A, and lysed by sonication at high power 3 times for 10 sec with one-minute interval. The lysates were centrifuged at 20,000 x g for 10 min at 4°C. The supernatant was incubated with 1 μg of rabbit anti-RASSF6 antibody or control rabbit IgG overnight at 4 °C, and was further incubated with protein G-Sepharose 4 fast-flow beads (GE Healthcare) for 2 h at 4 °C. The beads were washed four times with PBS. The precipitates were analyzed by SDS-PAGE and immunoblotting.

### Co-immunoprecipitation for exogenously expressed proteins

HEK293FT cells were plated at 1×10^6^ cells/well in a 6 well plate. 24 h later, the indicated plasmids were transfected with Lipofectamine 2000. 48 h later, the cells were treated with either 10 μM epoxomicin or 10 μM MG-132 for 6 h and then harvested. The cells were lysed in 500 μl of the lysis buffer (25 mM Tris-HCl pH7.4, 100 mM NaCl, 1 mM EDTA, 1 mM EGTA, 0.5% (w/v) sodium deoxycholate, 1%(v/v) TritonX-100, 50 μM APMSF, 3 mg/l pepstatin A, 10 mg/l leupeptin, 3 mg/l pepstatin A, 50 mM NaF, 2 mM Na_3_VO_4_, and 10 μM MG-132) and centrifuged at 20,000 x g for 10 min. The supernatant (the input) was incubated with 5 μl of anti-DYKDDDDK-tag beads, anti-V5-tag beads or anti-GFP antibody fixed on protein G-Sepharose 4 fast-flow beads. The beads were washed with the lysis buffer. The proteins in the inputs and in the immunoprecipitates were detected with antibodies. For Lumier assay, we used luciferase-fused proteins and measured luciferase activity in immunoprecipitates by use of Picagene (Toyo INK, Tokyo, Japan) as a substrate (56).

### Subcellular fractionation

Cells at 70-80% confluency in 60-mm dishes were washed with ice-cold PBS and harvested by scraping. Cells were collected by centrifugation at 4°C, resuspended with 200 μl of the hypotonic buffer (10mM Hepes-NaOH pH 7.5, 10 mM KCl, 1.5 mM MgCl_2_, 0.34 M sucrose, 10 % (v/v) glycerol, 0.05 % (v/v) Nonidet P-40, 50 mM APMSF, 10 mg/l leupeptin, 3 mg/l pepstatin, 50 mM NaF, 2 mM Na_3_VO_4_, and 25 mM β-glycerophosphate) and kept at 4°C for 5 min. Cells were then resuspended by pipetting. After 66.6μl of the mixture was saved as the whole cell lysate, the remaining samples were centrifuged at 800 x g at 4°C for 5 min. The supernatant was centrifuged again at 20,000 x g for 10 min at 4°C. 90μl of the supernatant was collected as the cytosolic fraction. The pellet was washed twice with the hypotonic buffer, suspended with 133.3 μl of the same buffer and used as the nuclear fraction.

### Monitoring active DNA synthesis

Active DNA synthesis was evaluated by use of Click-iT Edu Alexa 488 Imaging kit (#C10337) (Thermo Fisher Scientific) according to the manufacturer’s protocol.

### Apoptosis

GFP-RASSF6 proteins were expressed in HCT116 p53-/- cells. The cells were immunostained with anti-cytochrome C antibody and the nuclei were visualized with Hoechst 33342. The cytochrome C release and the nuclear condensation were evaluated. Detection of the sub-G1 population was performed by FACS as described previously (14).

### RNA interference

RNA interferences were performed by use of Lipofectamine RNAiMAX (Thermo Fisher Scientific). The following small interfering (si) RNAs are obtained from Thermo Fisher Scientific; Silencer Select negative control no.2; human RASSF6 #1 (s46640) and #2 (s46639); human E2F1 (s4405); human RB1 #1 (s523) and #2 (s522); human BMI1 (s2015); human LATS1 (s17392); human LATS2 (s25505); human RASSF1 (s22088); human RASSF5 (s38021); and human YAP1 #1 (s20366) and #2 (s20367). Human LIN52 (sc-92126) was obtained from Santa Cruz Biotechnology.

### Quantitative RT-PCR

Quantitative RT-PCR was performed in HCT116 p53-/- cells as described previously (14). Primers are listed in **Table 1**.

**Table 1.**
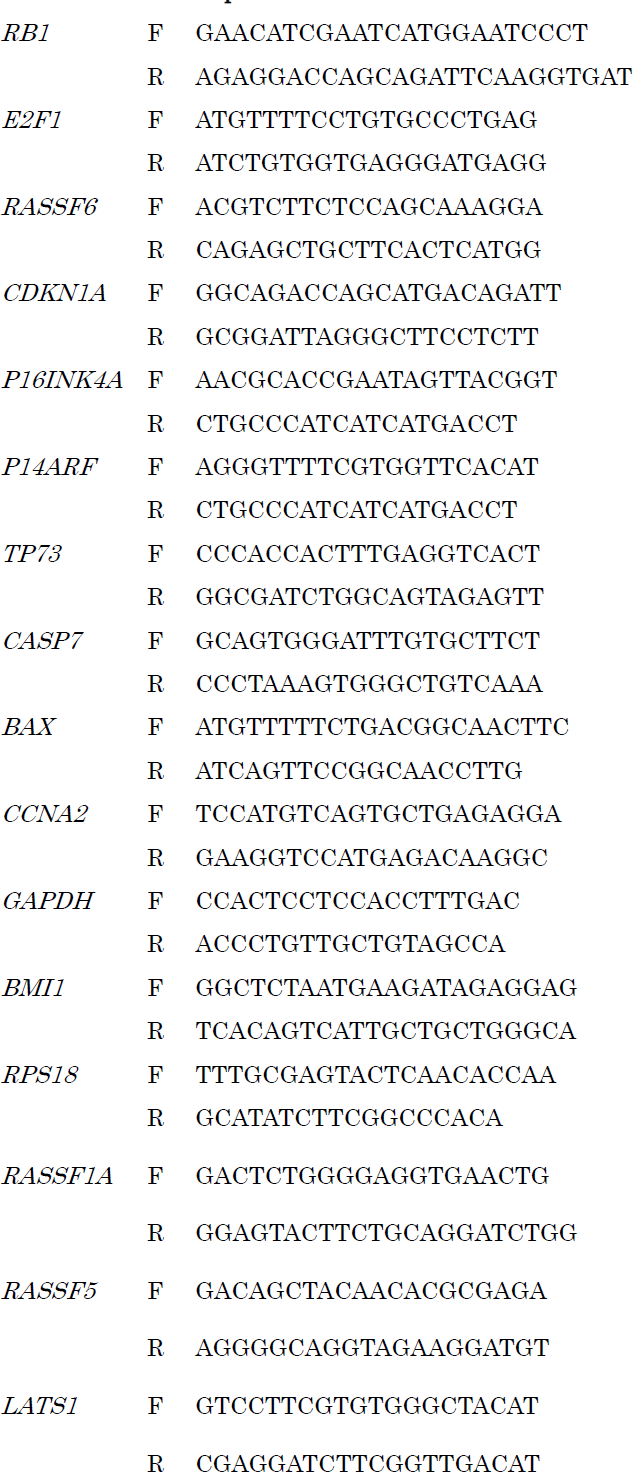
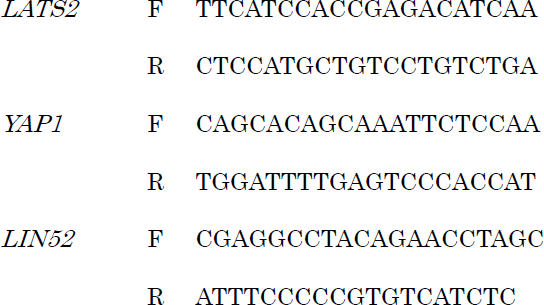
Primers for qRT-PCR

### Reporter assay

For the E2F1 reporter assay, HEK293FT cells were transfected with E2F1-Luc (−242) reporter (a gift of Masa-Aki Ikeda, Tokyo Medical and Dental University) and pCMV alkaline phosphatase (a gift of Sumiko Watanabe, The University of Tokyo). Luciferase activity was assayed as described previously (58).

### Treatment with lambda phosphatase

HCT116 p53-/- cells were plated at 4×105 cells in a 60-mm dish and transfected with control siRNA or *RASSF6* siRNA. 72 h later, the cells were treated with 50 mg/l cycloheximide for 3 h. Then the cells were collected and lysed in 200 μl of the above described lysis buffer. The lysates were centrifuged at 20,000 x g for 10 min. 39.5 μl of the supernatant was added by 200 unit (0.5 μl) of lambda protein phosphatase, 10 x lambda phosphatase buffer (500 mM Hepes-NaOH pH 7.5, 1 mM EGTA, 50 mM dithiothreitol and 0.1% Brij 35) and 5 μl of 20 mM MnCl2 and incubated for 30 mi at 30 °C.

### Ubiquitination of BMI1

HEK293FT cells were transfected with either control siRNA or *RASSF6*-targeted siRNA. 24 h later, the cells were further transfected with pCIneoFH-BMI1 and pCGN-HA-UBC. 48 h later, the cells were treated with 30 μM MG-132 for 6 h, harvested, and lysed in the denaturing buffer A (6 M guanidium hydrochloride, 100 mM Na_2_HPO_4_/NaH_2_PO_4_ pH8.0, 10 mM Tris-HCl pH8.0, and 10 Mm β-mercaptoethanol) by sonication. FH-BMI1 was isolated by use of Ni-NTA agarose beads (QIAGEN, Vento, Netherlands). The beads were washed with buffer B (8 M urea, 100 mM Na_2_HPO_4_/NaH_2_PO_4_ pH8.0, 10 mM Tris-HCl pH8.0, and 10 mM β-mercaptoethanol), FH-BMI1 was eluted with buffer C (200 mM Imidazole, 10 mM Tris-HCl pH6.7, 0.72 M β-mercaptoethanol, 5% (w/v) SDS, and 30% glycerol). The eluents were analyzed by the immunoblotting with anti-HA and anti-FLAG antibodies.

### Statistical Analysis

Statistical analyses were performed with Student’s *t* test for comparison between two samples and analysis of variance with Bonferroni’s post hoc test for multiple comparisons using GraphPad Prism software (GraphPad Software).

## ACKNOWLEDGEMENTS

We are grateful for Dr. Masa-Aki Ikeda (Tokyo Medical and Dental University), Dr. Akira Kikuchi (Osaka University), Dr. Yue Xiong (University of North Carolina) for the materials. Plasmid collections of lentiviral human cDNA expression library (GE Dharmacon, UK) were supplied from TMDU Gene Library. S.H. and A.S. are supported by the MEXT scholarship. This work was supported by research grants from Japan Society for the Promotion of Science (JSPS) (26460359, 26293061), and the Mitsubishi Foundation (26138). The authors declare no conflict of interest.

## REFERENCES

1. Avruch J, Zhou D, Fitamant J, Bardeesy N, Mou F, Barrufet LR. 2012. Protein kinases of the Hippo pathway: regulation and substrates. Semin Cell Dev Biol 23:770–784.

2. Volodko N, Gordon M, Salla M, Ghazaleh HA, Baksh S. 2014. RASSF tumor suppressor gene family: Biological functions and regulation. FEBS Lett.

3. Iwasa H, Hossain S, Hata Y. 2018. Tumor suppressor C-RASSF proteins. Cell Mol Life Sci.

4. Hesson LB, Dunwell TL, Cooper WN, Catchpoole D, Brini AT, Chiaramonte R, Griffiths M, Chalmers AD, Maher ER, Latif F. 2009. The novel RASSF6 and RASSF10 candidate tumour suppressor genes are frequently epigenetically inactivated in childhood leukaemias. Mol Cancer 8:42.

5. Shinawi T, Hill V, Dagklis A, Baliakas P, Stamatopoulos K, Agathanggelou A, Stankovic T, Maher ER, Ghia P, Latif F. 2012. KIBRA gene methylation is associated with unfavorable biological prognostic parameters in chronic lymphocytic leukemia. Epigenetics 7:211–215.

6. Djos A, Martinsson T, Kogner P, Carén H. 2012. The RASSF gene family members RASSF5, RASSF6 and RASSF7 show frequent DNA methylation in neuroblastoma. Mol Cancer 11:40.

7. Mezzanotte JJ, Hill V, Schmidt ML, Shinawi T, Tommasi S, Krex D, Schackert G, Pfeifer GP, Latif F, Clark GJ. 2014. RASSF6 exhibits promoter hypermethylation in metastatic melanoma and inhibits invasion in melanoma cells. Epigenetics 9:1496– 1503.

8. Guo W, Dong Z, Guo Y, Shen S, Guo X, Kuang G, Yang Z. 2015. Decreased expression and frequent promoter hypermethylation of RASSF2 and RASSF6 correlate with malignant progression and poor prognosis of gastric cardia adenocarcinoma. Mol Carcinog.

9. Wen Y, Wang Q, Zhou C, Yan D, Qiu G, Yang C, Tang H, Peng Z. 2011. Decreased expression of RASSF6 is a novel independent prognostic marker of a worse outcome in gastric cancer patients after curative surgery. Ann Surg Oncol 18:3858–3867.

10. Ye HL, Li DD, Lin Q, Zhou Y, Zhou QB, Zeng B, Fu ZQ, Gao WC, Liu YM, Chen RW, Li ZH, Chen RF. 2015. Low RASSF6 expression in pancreatic ductal adenocarcinoma is associated with poor survival. World J Gastroenterol 21:6621–6630.

11. Ikeda M, Hirabayashi S, Fujiwara N, Mori H, Kawata A, Iida J, Bao Y, Sato Y, Iida T, Sugimura H, Hata Y. 2007. Ras-association domain family protein 6 induces apoptosis via both caspase-dependent and caspase-independent pathways. Exp Cell Res 313:1484–1495.

12. Ikeda M, Kawata A, Nishikawa M, Tateishi Y, Yamaguchi M, Nakagawa K, Hirabayashi S, Bao Y, Hidaka S, Hirata Y, Hata Y. 2009. Hippo pathway-dependent and -independent roles of RASSF6. Sci Signal 2:ra59.

13. Withanage K, Nakagawa K, Ikeda M, Kurihara H, Kudo T, Yang Z, Sakane A, Sasaki T, Hata Y. 2012. Expression of RASSF6 in kidney and the implication of RASSF6 and the Hippo pathway in the sorbitol-induced apoptosis in renal proximal tubular epithelial cells. J Biochem 152:111–119.

14. Iwasa H, Kudo T, Maimaiti S, Ikeda M, Maruyama J, Nakagawa K, Hata Y. 2013. The RASSF6 tumor suppressor protein regulates apoptosis and the cell cycle via MDM2 protein and p53 protein. J Biol Chem 288:30320–30329.

15. Pan D. 2010. The hippo signaling pathway in development and cancer. Dev Cell 19:491–505.

16. Kodaka M, Hata Y. 2014. The mammalian Hippo pathway: regulation and function of YAP1 and TAZ. Cell Mol Life Sci.

17. Meng Z, Moroishi T, Guan KL. 2016. Mechanisms of Hippo pathway regulation. Genes Dev 30:1–17.

18. Allen NP, Donninger H, Vos MD, Eckfeld K, Hesson L, Gordon L, Birrer MJ, Latif F, Clark GJ. 2007. RASSF6 is a novel member of the RASSF family of tumor suppressors. Oncogene 26:6203–6211.

19. Sarkar A, Iwasa H, Hossain S, Xu X, Sawada T, Shimizu T, Maruyama J, Arimoto-Matsuzaki K, Hata Y. 2017. Domain analysis of Ras-association domain family member 6 upon interaction with MDM2. FEBS Lett 591:260–272.

20. Weinberg RA. 1995. The retinoblastoma protein and cell cycle control. Cell 81:323– 330.

21. Vogelstein B, Lane D, Levine AJ. 2000. Surfing the p53 network. Nature 408:307–310.

22. Vousden KH, Lane DP. 2007. p53 in health and disease. Nat Rev Mol Cell Biol 8:275– 283.

23. Dyson NJ. 2016. RB1: a prototype tumor suppressor and an enigma. Genes Dev 30:1492–1502.

24. Rubin SM. 2013. Deciphering the retinoblastoma protein phosphorylation code. Trends Biochem Sci 38:12–19.

25. Macdonald JI, Dick FA. 2012. Posttranslational modifications of the retinoblastoma tumor suppressor protein as determinants of function. Genes Cancer 3:619–633.

26. Dick FA, Rubin SM. 2013. Molecular mechanisms underlying RB protein function. Nat Rev Mol Cell Biol 14:297–306.

27. Rubin SM, Gall AL, Zheng N, Pavletich NP. 2005. Structure of the Rb C-terminal domain bound to E2F1-DP1: a mechanism for phosphorylation-induced E2F release. Cell 123:1093–1106.

28. Burke JR, Deshong AJ, Pelton JG, Rubin SM. 2010. Phosphorylation-induced conformational changes in the retinoblastoma protein inhibit E2F transactivation domain binding. J Biol Chem 285:16286–16293.

29. Zarkowska T, Mittnacht S. 1997. Differential phosphorylation of the retinoblastoma protein by G1/S cyclin-dependent kinases. J Biol Chem 272:12738–12746.

30. Lundberg AS, Weinberg RA. 1998. Functional inactivation of the retinoblastoma protein requires sequential modification by at least two distinct cyclin-cdk complexes. Mol Cell Biol 18:753–761.

31. Ezhevsky SA, Nagahara H, Vocero-Akbani AM, Gius DR, Wei MC, Dowdy SF. 1997. Hypo-phosphorylation of the retinoblastoma protein (pRb) by cyclin D:Cdk4/6 complexes results in active pRb. Proc Natl Acad Sci U S A 94:10699–10704.

32. Antonucci LA, Egger JV, Krucher NA. 2014. Phosphorylation of the Retinoblastoma protein (Rb) on serine-807 is required for association with Bax. Cell Cycle 13:3611– 3617.

33. Ren S, Rollins BJ. 2004. Cyclin C/cdk3 promotes Rb-dependent G0 exit. Cell 117:239– 251.

34. Tschöp K, Conery AR, Litovchick L, Decaprio JA, Settleman J, Harlow E, Dyson N. 2011. A kinase shRNA screen links LATS2 and the pRB tumor suppressor. Genes Dev 25:814–830.

35. Kolupaeva V, Janssens V. 2013. PP1 and PP2A phosphatases--cooperating partners in modulating retinoblastoma protein activation. FEBS J 280:627–643.

36. Ribeiro PS, Josué F, Wepf A, Wehr MC, Rinner O, Kelly G, Tapon N, Gstaiger M. 2010. Combined functional genomic and proteomic approaches identify a PP2A complex as a negative regulator of Hippo signaling. Mol Cell 39:521–534.

37. Shi Z, Jiao S, Zhou Z. 2016. STRIPAK complexes in cell signaling and cancer. Oncogene.

38. Couzens AL, Knight JD, Kean MJ, Teo G, Weiss A, Dunham WH, Lin ZY, Bagshaw RD, Sicheri F, Pawson T, Wrana JL, Choi H, Gingras AC. 2013. Protein interaction network of the Mammalian hippo pathway reveals mechanisms of kinase-phosphatase interactions. Sci Signal 6:rs15.

39. Barnoud T, Donninger H, Clark GJ. 2016. Ras Regulates Rb via NORE1A. J Biol Chem 291:3114–3123.

40. Guo C, Zhang X, Pfeifer GP. 2011. The tumor suppressor RASSF1A prevents dephosphorylation of the mammalian STE20-like kinases MST1 and MST2. J Biol Chem 286:6253–6261.

41. Iwasa H, Jiang X, Hata Y. 2015. RASSF6; the Putative Tumor Suppressor of the RASSF Family. Cancers (Basel) 7:2415–2426.

42. Uchida C, Miwa S, Kitagawa K, Hattori T, Isobe T, Otani S, Oda T, Sugimura H, Kamijo T, Ookawa K, Yasuda H, Kitagawa M. 2005. Enhanced Mdm2 activity inhibits pRB function via ubiquitin-dependent degradation. EMBO J 24:160–169.

43. Jacobs JJ, Kieboom K, Marino S, DePinho RA, van Lohuizen M. 1999. The oncogene and Polycomb-group gene bmi-1 regulates cell proliferation and senescence through the ink4a locus. Nature 397:164–168.

44. Sahasrabuddhe AA, Dimri M, Bommi PV, Dimri GP. 2011. βTrCP regulates BMI1 protein turnover via ubiquitination and degradation. Cell Cycle 10:1322–1330.

45. Indovina P, Pentimalli F, Casini N, Vocca I, Giordano A. 2015. RB1 dual role in proliferation and apoptosis: cell fate control and implications for cancer therapy. Oncotarget 6:17873–17890.

46. Day ML, Foster RG, Day KC, Zhao X, Humphrey P, Swanson P, Postigo AA, Zhang SH, Dean DC. 1997. Cell anchorage regulates apoptosis through the retinoblastoma tumor suppressor/E2F pathway. J Biol Chem 272:8125–8128.

47. Ianari A, Natale T, Calo E, Ferretti E, Alesse E, Screpanti I, Haigis K, Gulino A, Lees JA. 2009. Proapoptotic function of the retinoblastoma tumor suppressor protein. Cancer Cell 15:184–194.

48. Müller H, Bracken AP, Vernell R, Moroni MC, Christians F, Grassilli E, Prosperini E, Vigo E, Oliner JD, Helin K. 2001. E2Fs regulate the expression of genes involved in differentiation, development, proliferation, and apoptosis. Genes Dev 15:267–285.

49. Moroni MC, Hickman ES, Lazzerini Denchi E, Caprara G, Colli E, Cecconi F, Müller H, Helin K. 2001. Apaf-1 is a transcriptional target for E2F and p53. Nat Cell Biol 3:552–558.

50. Nahle Z, Polakoff J, Davuluri RV, McCurrach ME, Jacobson MD, Narita M, Zhang MQ, Lazebnik Y, Bar-Sagi D, Lowe SW. 2002. Direct coupling of the cell cycle and cell death machinery by E2F. Nat Cell Biol 4:859–864.

51. Korah J, Falah N, Lacerte A, Lebrun JJ. 2012. A transcriptionally active pRb-E2F1-P/CAF signaling pathway is central to TGFβ-mediated apoptosis. Cell Death Dis 3:e407.

52. Pediconi N, Ianari A, Costanzo A, Belloni L, Gallo R, Cimino L, Porcellini A, Screpanti I, Balsano C, Alesse E, Gulino A, Levrero M. 2003. Differential regulation of E2F1 apoptotic target genes in response to DNA damage. Nat Cell Biol 5:552–558.

53. Banerjee Mustafi S, Chakraborty PK, Dwivedi SK, Ding K, Moxley KM, Mukherjee P, Bhattacharya R. 2017. BMI1, a new target of CK2α. Mol Cancer 16:56.

54. Ma S, Tang J, Feng J, Xu Y, Yu X, Deng Q, Lu Y. 2008. Induction of p21 by p65 in p53 null cells treated with Doxorubicin. Biochim Biophys Acta 1783:935–940.

55. Vayssade M, Haddada H, Faridoni-Laurens L, Tourpin S, Valent A, Bénard J, Ahomadegbe JC. 2005. P73 functionally replaces p53 in Adriamycin-treated, p53-deficient breast cancer cells. Int J Cancer 116:860–869.

56. Kawano S, Maruyama J, Nagashima S, Inami K, Qiu W, Iwasa H, Nakagawa K, Ishigami-Yuasa M, Kagechika H, Nishina H, Hata Y. 2015. A cell-based screening for TAZ activators identifies ethacridine, a widely used antiseptic and abortifacient, as a compound that promotes dephosphorylation of TAZ and inhibits adipogenesis in C3H10T1/2 cells. J Biochem 158:413–423.

57. Ohta T, Xiong Y. 2001. Phosphorylation- and Skp1-independent in vitro ubiquitination of E2F1 by multiple ROC-cullin ligases. Cancer Res 61:1347–1353.

58. Bao Y, Nakagawa K, Yang Z, Ikeda M, Withanage K, Ishigami-Yuasa M, Okuno Y, Hata S, Nishina H, Hata Y. 2011. A cell-based assay to screen stimulators of the Hippo pathway reveals the inhibitory effect of dobutamine on the YAP-dependent gene transcription. J Biochem 150:199–208.

